# Temporal emergence of age-associated changes in cognitive and physical function in vervets (*Chlorocebus aethiops sabaeus*)

**DOI:** 10.1101/2020.09.28.313460

**Authors:** Brett M. Frye, Payton M. Valure, Suzanne Craft, Mark G. Baxter, Christie Scott, Shanna Wise-Walden, David Bissinger, Hannah M. Register, Carson Copeland, Matthew J. Jorgensen, Jamie N. Justice, Stephen B. Kritchevsky, Thomas C. Register, Carol A. Shively

## Abstract

Dual declines in gait speed and cognitive performance are associated with increased risk of developing dementia. Characterizing the patterns of such impairments therefore is paramount to distinguishing healthy from pathological aging. Nonhuman primates such as vervet/African green monkeys (*Chlorocebus aethiops sabaeus*) are important models of human neurocognitive aging, yet the trajectory of dual decline has not been characterized. We therefore 1) assessed whether cognitive and physical performance (i.e., gait speed) are lower in older aged animals; 2) explored the relationship between performance in a novel task of executive function (Wake Forest Maze Task – WFMT) and a well-established assessment of working memory (Delayed Response Task – DR Task); and 3) examined the association between baseline gait speed with executive function and working memory at one-year follow-up. We found 1) physical and cognitive declines with age; 2) strong agreement between performance in the novel WFMT and DR task; and 3) that slow gait predicted poor cognitive performance in both domains. Our results suggest that older-aged vervets exhibit a coordinated suite of traits consistent with human aging and that slow gait may be a risk factor for cognitive decline. This integrative approach provides evidence that gait speed and cognitive function differ across the lifespan in female vervet monkeys, which advances them as a model that could be used to evaluate the trajectory of dual decline over time.

## INTRODUCTION

The average life expectancy in the United States has increased from 70 to nearly 79 years over the last half century [1]. However, longer lives are not always accompanied by years of healthy, active life [2]. Slow gait increasingly is recognized as an early biomarker for poor aging outcomes, with slower-than-average gaits being linked to increased risk for cardiovascular disease [3] and early mortality [4, 5]. Mounting evidence also has shown direct associations between gait and cognitive performance, suggesting common underlying processes [6]. Dual declines in physical and cognitive performance may be particularly problematic for neurocognitive health [7–9], as individuals exhibiting both cognitive and physical impairments are three to 6.28 times more likely to develop dementia than are individuals without these phenotypes [10–12].

Characterizing the temporal patterns of mobility and cognitive declines may aid early detection of pathological aging, yet the trajectories of such age-related declines remains an area of inquiry (e.g., [13]). Cross-sectional studies have demonstrated that impaired cognitive performance predicts deficits in physical capabilities [14, 8]. These studies suggest that progressive cognitive dysfunction promotes functional impairments. Others have reported that functional impairments precede declines in cognitive capabilities [15–20]. If so, physical function would serve as an early biomarker for neurocognitive risk. In addition to inconsistent findings about the temporal patterns of emergence, the evidence linking physical performance to particular cognitive domains is mixed, with varying degrees of association being drawn between executive function and working memory [21]. Thus, additional study is needed to 1) explore the age-associated declines in both physical performance and cognitive function, and 2) examine the strength of associations between physical performance characteristics and cognitive domain phenotypes.

The clinical cohort studies needed to study aging processes can be difficult and expensive. Preclinical animal studies may provide alternative, timely, and cost-effective insights into aging phenotypes across the life course [22]. Nonhuman primates (NHPs) are particularly well-poised for such investigations as they recapitulate many brain and body changes observed in human aging [23]. Vervets, also known as African green monkeys (*Chlorocebus aethiops sabaeus*), are among the best characterized NHP models of chronic disease risk and age-related conditions [24–26]. Additionally, vervets are increasingly used to study neuropathologic changes associated with Alzheimer’s disease (AD) [27, 28]. Older-aged vervets develop Aβ plaques naturally with age in patterns similar to humans with AD. Importantly, greater densities of Aβ plaques are associated with slower gait speeds, which suggests that slow gait speed may represent a translationally relevant biomarker of neuropathologic aging [27].

Considering the strengths of the vervet model, characterization of their use as a translational model of functional aging is greatly needed. We therefore investigated cognitive and physical performance (i.e., gait speed) in an aging cohort of adult female vervets (n=30; 8 to 29 years old). Our aims were threefold: 1) to assess whether cognitive performance and gait speed are lower in older aged animals; 2) to explore the relationship between a novel task of executive function – the Wake Forest Maze Task (WFMT) – and a well-established assessment of working memory; and 3) to examine the association between baseline gait speed with executive function and working memory at one-year follow-up. We hypothesized that cognitive and physical performance would be lower in older aged animals. We also predicted that performance in the novel executive function task would reflect that in the conventional assessment of working memory. Lastly, we expected that slower gait speed would predict poorer performance in both working memory and executive function at one-year follow-up.

## METHODS

### Subjects & Housing

Our sample consisted of 30 socially housed female vervet monkeys (*Chlorocebus aethiops sabaeus*) ranging in age from 8 to 29 years old. Three age-classes characterized the cohort: middle-aged (N=10; 8-15 years old), older (N=6; 16-20 years old), and oldest (N=14; 21-29 years old) (Subject characteristics; **Supplementary Table 1**). Study subjects were born and raised into the Vervet Research Colony (VRC) (Wake Forest School of Medicine). The VRC is a breeding colony made up of approximately 300 animals living in social groups that reflect similar age and sex-class compositions as in naturalistic settings. This setting has been designed to optimize studies related to development, aging, and chronic disease risk across the life span. Animals’ routine diets consisted of LabDiet Monkey Chow. During cognitive assessments, we delayed feeding until after assessment to standardize motivation to participate in the tasks. Water was available *ad libitum* prior to and after testing. All observations complied with state and federal regulations and with the approval of the Wake Forest School of Medicine institutional animal care and use committee.

### Experimental Design

Vervets’ enclosures consisted of a large outdoor area (34×30’) connected to an indoor enclosure (10×30’) via capture tunnel access. Data of gait speed were collected in the outdoor portions of the enclosures. Cognitive assessments were conducted indoors, using apparatus that could be affixed to the end of the capture tunnels. Social partners of the focal monkey were kept outside during cognitive tests such that the focal maintained auditory but not visual contact with their group. Thus, monkeys were tested individually within their home enclosures but without interference from social partners. All cognitive assessments were video recorded. Gait speeds were collected approximately one-year before cognitive performance testing in order to assess whether gait speed predicted later executive function and working memory capabilities.

### Physical Performance – Gait Speed

Gait speed represented the biomarker of physical performance and was collected using a well-validated comparative assessment; we quantified gait speeds using a stopwatch to measure the time it took an individual to traverse 3 feet at a normal, unprovoked (i.e., not fleeing or chasing) pace [29, 30]. We measured gait speeds during the day (0900-1600) and calculated average gait speeds (time per distance traversed (cm/s)) using five instances.

### Cognitive Performance – Wake Forest Maze Task

#### Apparatus

We adapted a puzzle feeder apparatus (Primate Products, Immokalee, FL) to develop the Wake Forest Maze Task (WFMT) to assess executive function. This procedure was modified from previous methods of assessing age-related declines in cynomolgus monkeys (*Macaca fascicularis*) [31]. As previously described by Watson et al. (1999), the puzzle feeder consisted of a Plexiglas box (24.13 cm h × 22.23 cm w × 4.45 cm d) into which horizontal and vertical tabs could be inserted to generate maze configurations of varying levels of difficulty. The side of the puzzle facing the monkey had small slits that allowed the animal to manually manipulate, but not retrieve, food items within the box. We used food items (baby carrots, halved) that were durable enough that they could only be retrieved from enlarged openings at the base of the puzzle, after the animal had successfully moved the food item through the maze. Food rewards included baby carrots (during testing) and grapes (during acclimation and as positive reinforcement training).

#### Acclimation

Monkeys in this study did not have prior experience with the puzzle feeder apparatus. Thus, each vervet underwent an acclimation period lasting no longer thirty minutes per day, over the course of five days. Briefly, this acclimation procedure involved 1) enticing the monkey to interact with the puzzle feeder by loading a randomly assorted maze with high quality food rewards (grapes), 2) replacing high quality foods with less desirable (but more durable) food items (carrots) followed by positive reinforcement (i.e., clicker mark followed by a high-quality reward), and 3) practice in moving a food reward over a vertical insert (**Supplementary Fig. 1**). The last stage of acclimation was added after pilot data demonstrated that monkeys could not perform the maze task without this prior training. This stage also allowed us to assess whether subjects had sufficient manual dexterity to complete all levels of the task.

#### Testing

The maze task was presented to individuals immediately following the acclimation period. Testing consisted of 30-minute sessions over five consecutive days. During each session, the vervet manipulated the low-quality food reward through a series of 19 increasingly difficult maze configurations (**Supplementary Fig. 2**). If the vervet successfully maneuvered the food item to the large opening at the bottom of the maze, she immediately received a mark (i.e., clicker noise) and high-quality reward (grape). In the least difficult levels (Levels 1-6), the animals only had to manipulate objects horizontally. At higher levels, animals were also required to maneuver the food item vertically. The maze was only configured to increase the level of difficulty when the previous level had been solved. This design allowed animals to progress through as many levels as possible during the 30-minute session.

Some animals exhibited a directional bias: i.e., monkeys readily interacted with the apparatus if the initial movement was in either the left or right direction but would otherwise would not conduct the maze task. Thus, we developed mirror images – i.e., a left and a right series – for each maze level. Animals were allowed to progress though either the left or the right series continuously. Sometimes, the animals ceased to interact with the maze. When this occurred, testing was paused, and the animal was presented with the maze arranged in the same manner as during the acclimation period. Upon successful retrieval from the maze, we positively reinforced the behavior. This step was sometimes necessary to entice the monkeys to interact with the apparatus. After this, we resumed normal testing. The time it took the monkey to complete this “reset” task was not deducted from that session’s 30-minutes time limit.

### Cognitive Performance – Working Memory Task of Delayed Response

#### Apparatus

We used a modified Wisconsin General Test apparatus to assess working memory via a task of delayed response (DR) (**Supplemental Fig. 3**). This DR task was adapted from previous assessments in nonhuman primates [32–34]. Here, the apparatus had three equidistant reward boxes and the screen separating the animal from the reward boxes was transparent instead of opaque, as pilot data revealed that animals avoided opaque screens and would not subsequently participate in the task.

#### Training

We trained vervets to access the reward boxes by baiting each of the three identical reward boxes with a high-quality food reward (grapes, quartered). During this phase of the experiment, the lids on the equidistant boxes remained open such that the monkey could clearly see the reward. We repeated this process for three trials. Next, we baited all three reward boxes and closed the lids. This step ensured that the monkeys had sufficient dexterity to retrieve the rewards. We repeated this process another three times. After successfully demonstrating ability to open each of the reward boxes, subjects advanced to a series of trials in which a single reward box contained a food reward. For this and all subsequent stages of the assessment, trials began with the reward boxes open and the transparent partition lowered. This ensured the monkey could see which reward box was baited by could not access the reward (procedure comparable to [35]).

To begin, we baited a single reward box and then covered the boxes with lids, closing the baited box last. Immediately after closing the baited box, we raised the transparent screen (1 second delay) to permit a response. The subject was permitted to select (i.e., touch) a single box. If she selected the baited box, the subject was allowed to retrieve the reward. This constituted a “correct” response. If the subject selected a non-baited box, the partition was immediately lowered, and an “incorrect” score was recorded. We repeated this process for thirty trials. The reward location was randomized. Monkeys received this training until they received a criterion of 80% success (≤6 errors during 30 trials) over three consecutive days.

#### Testing

Testing with increasingly longer delays followed training using an identical method to the 1 second delay. We introduced delays of 2.5, 5, 10, 15, 20 second delays. We tested each delay twice, so that each animal conducted a total of 60 trials at each delay. Some animals easily completed the 20 second delay. Animals that performed better than chance (33%) at the 20 second delay, advanced to 30, then 40, and then 60 second delays to the protocol. Performance greater than 33% at each delay prompted the subject to advance to these longer delays.

### Data Analysis

Data are presented as mean ± SEM. Statistical analysis was performed with the R version 4.0.0 (R Core Team, 2020). A value of p<0.05 was considered statistically significant. We inspected all data for outliers and visually inspected the distributions for normality. Where data deviated from normality, we analyzed both the transformed and non-transformed data. If transformation did not impact the outcome, we report the results of analyses of non-transformed data.

#### Wake Forest Maze Task (WFMT)

In our tests of executive function, we used both linear regression and one-way ANOVA to assess performance by age in the WFMT. Dependent variables were the highest level completed on either the left or right series, the total number of levels completed as sum of completed levels in both left and right series, and the total time spent interacting with the maze (i.e., manipulating food items).

#### Delayed Response (DR) Task

In our assessments of working memory, we used both linear regression and one-way ANOVA to assess performance by age in rate of acquisition of the DR task (i.e., trials to 80% criterion). We used a Pearson’s Chi-square test to assess whether age-groups differed in their propensities to reach the longest delay (60 seconds). We used mixed-effects logistic regression to analyze the proportions of trials in which individuals responded correctly during the DR task, after having acquired the task at the one-second delay. In these models, we tested for main- and interaction effects of three variables: age group (middle-aged, older, and oldest), time point (first or second attempt at given delay) and the delay (length of delay in seconds: 2.5-60 seconds). We included a random intercept term of individual to account for non-independence of repeated measures.

#### Relationship between WFMT & DR Task

We examined levels of agreement between the WFMT and DR tests with a series of Pearson’s correlations between the highest and sum of levels completed in the WFMT and the average accuracy at each delay.

#### Gait Speed and Cognitive Function at 1-year

We used linear regression and one-way ANOVA to assess whether gait speed predicted cognitive performance in both cognitive domains. For the WFMT, dependent variables were the highest (either left or right series) and total levels completed. For the DR task, the dependent variable was the average number of correct trials at the 20-second delay, i.e., the longest delay at which all subjects were tested.

## RESULTS

Gait speed data was collected for all study subjects (N=30). We used only three instances of walking to generate average gait speed for one middle-aged monkey because she had given birth and was actively carrying the infant for the remainder of the study period. WFMT was completed in 29 animals; one of the oldest monkeys could not complete task due to impaired dexterity. DR task was completed in 27 animals; three animals died of natural causes before completing the DR task.

### Age Effects on Cognitive Performance

#### Wake Forest Maze Task (WFMT)

We analyzed performance via the highest maze level completed and the total number of levels. Linear regression indicated that performance in the WFMT – assessed as highest level completed in either the left or right series – tended to decline with age, but the relationship was not statistically significant (t=-1.451, p=0.158) (**Fig. 1A**). Age was a significant predictor of the total number of maze levels achieved (t=-2.539, p=0.017) (**Fig. 1C**). One-way ANOVAs showed significant differences between the age groups in both the highest (F2,26)=4.915, p=0.015) (**Fig. 1B**) and total number of levels completed (F(2,26)=5.917; p-0.008) (**Fig. 1D**). *Post hoc* pairwise comparisons (p values adjusted using Tukey method for comparing a family of 3 estimates) indicated that animals in the older age class (16-20 years old) completed higher (more difficult) maze levels than did the oldest animals (p= 0.014). *Post hoc* comparisons also showed that middle-aged (p=0.022) and older vervets (p=0.023) completed more levels overall than did individuals within the oldest group. Age was unrelated to the total time spent interacting with the maze (regression: t=-1.31, p=0.202; ANOVA F(1,26)=1.911, p=0.168).

**Fig. 1.**
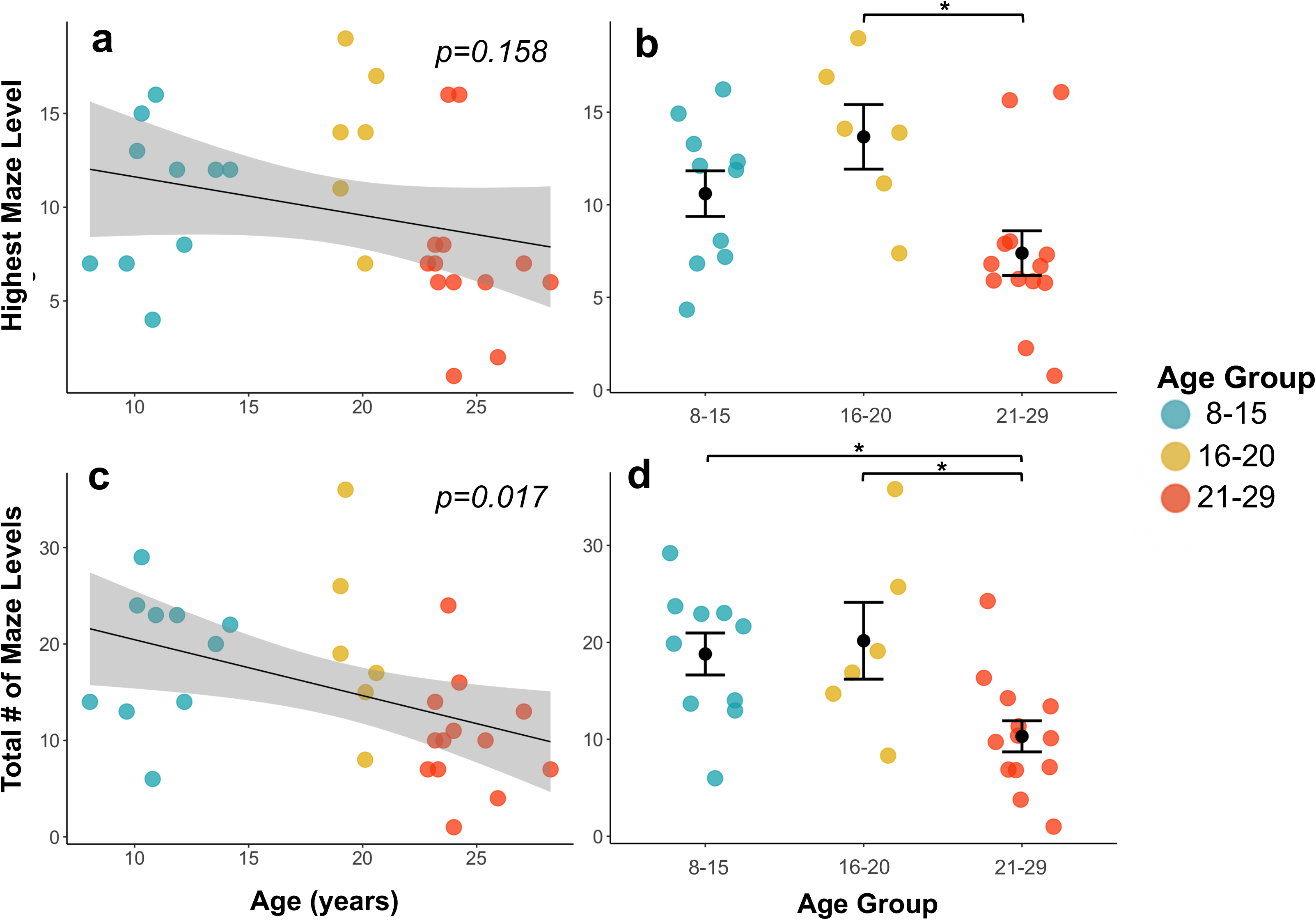
Age-based performance on the Wake Forest Maze Task (WFMT). a. The highest maze level completed (on either the left or right series) tended to decline with age (linear regression: t=-1.451, p=0.158); b. Older monkeys achieved higher levels on either the left or right maze series in the WFMT than did the oldest animals (ANOVA: F(2,26)=4.915; Tukey p=0.014); c. Total overall maze levels completed in the WFMT declined with age (linear regression: t=-2.539, p=0.017); d. Middle-aged (Tukey p=0.022) and older (Tukey p=0.023) monkeys completed more levels than did the oldest vervets (ANOVA: F(2,26)=5.917) (N=29) *p<0.05

#### Delayed Response Task

Age did not impact the rate with which animals acquired the DR task (i.e., number of trials to 80% criterion) when analyzed as either a continuous (regression: t=1.382, p=0.792) or categorical variable (ANOVA: (F(2,24)=1.644, p=0.214). Age groups did differ in their propensity to reach the longest (most difficult) delay of 60 seconds (χ^2^ =10.177, df=2, p=0.006). All middle-aged and older vervets reached the longest delays, whereas the majority of oldest vervets failed to reach the longest, i.e., most difficult, delay (**Supplementary Fig. 4**). Performance across delays, ranging from 2.5 to 60 seconds, for each age group is illustrated in **Fig. 2** as the percentage of correct trials. Logistic regression revealed that the effect of delay significantly interacted with age group (Likelihood Ratio Test: χ^2^=76.124, df=14, p<0.001). *Post-hoc* Tukey test comparisons of age groups at each delay showed significant divergence in performance at the 15, 20, 30, and 40 second delays, with the oldest vervets performing the more poorly than middle-aged and older vervets (**Table 1**).

**Fig. 2.**
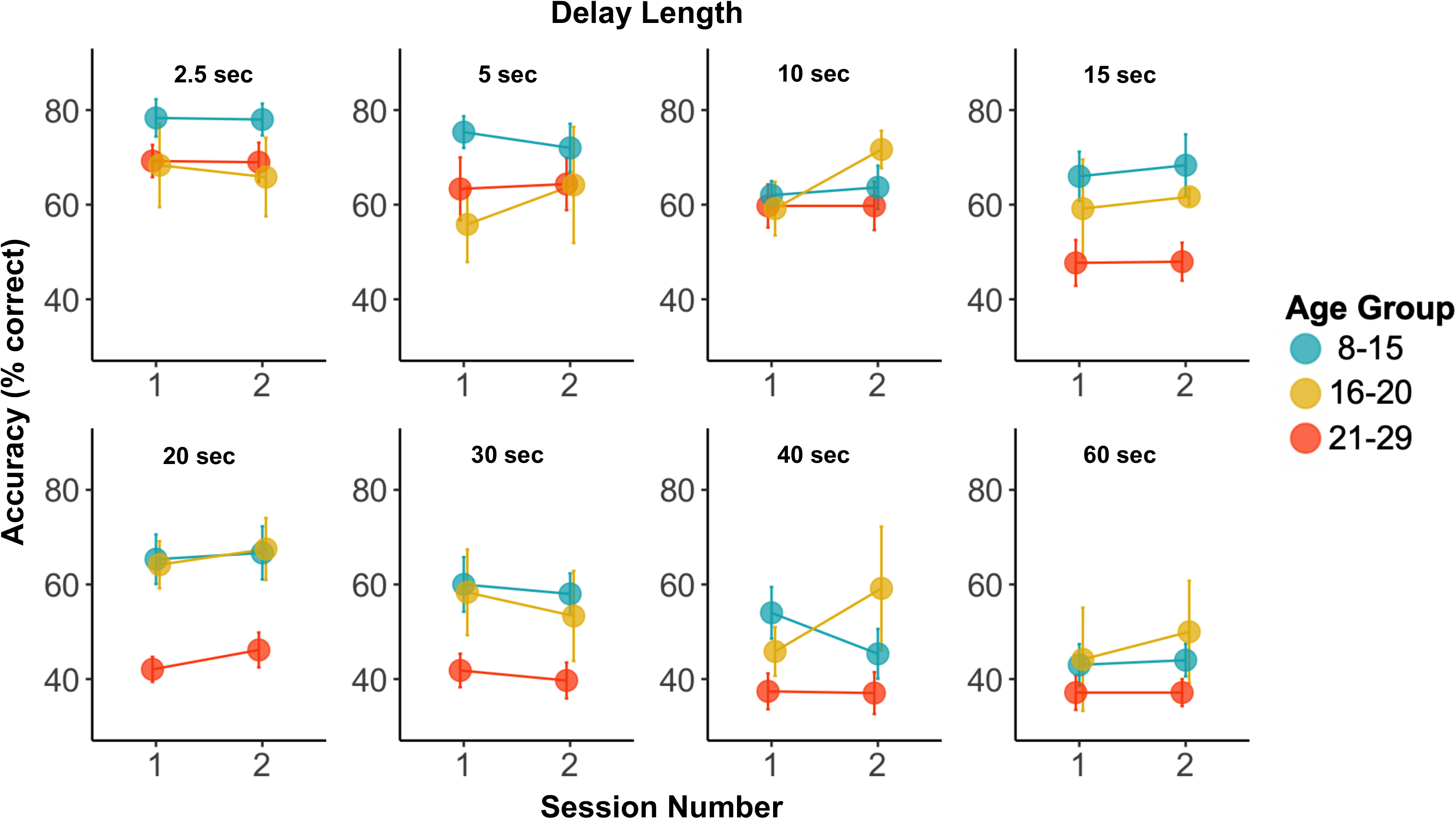
Accuracy (i.e., percent correct) across delays, ranging from 2.5 to 60 second, in the Delayed Response (DR) task for working memory. Logistic regression indicated that the oldest vervets (21-29 years old) performed more poorly than did the other age groups at the 15, 20, 30, and 40 second delays (N=27)

**Table 1.**
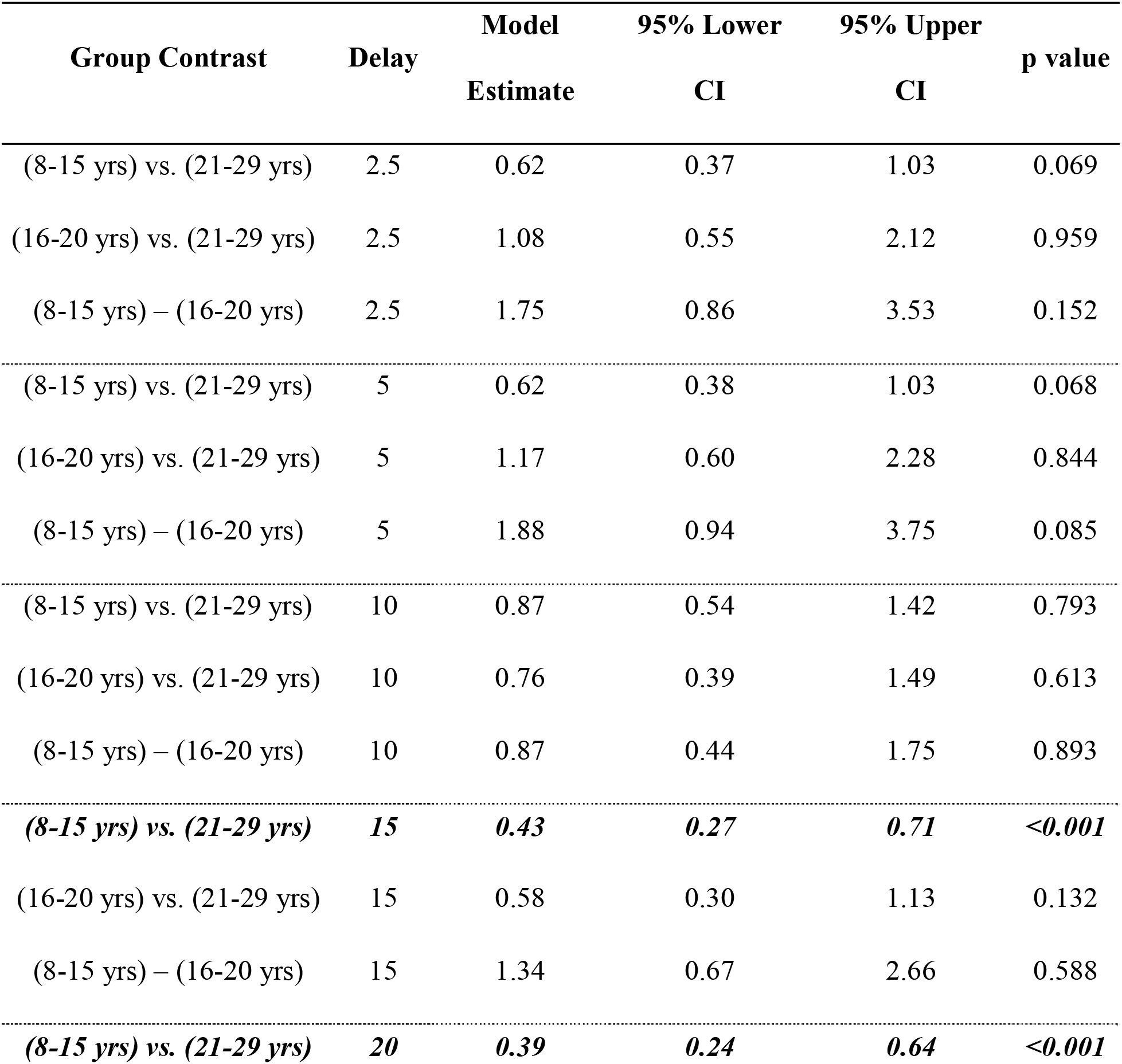
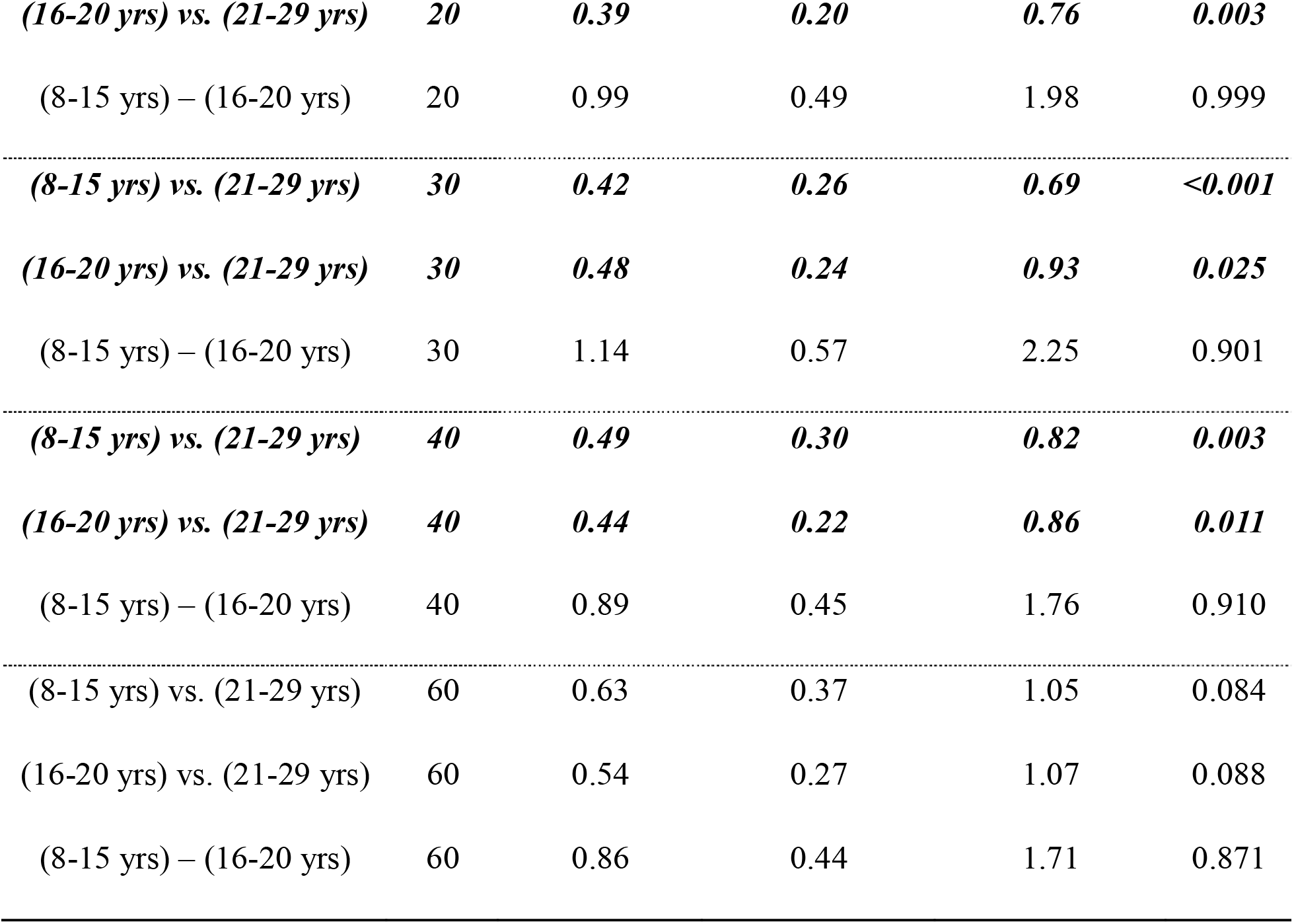
Post-hoc Tukey test comparisons of age groups at each delay showed significant divergence in performance at the 15, 20, 30, and 40 second delays. The oldest animals performed the more poorly than the youngest animals at the 15, 20, 30, and 40 second delays. Those in the oldest age group also performed more poorly than the 16-20 year old animals at the 20, 30, and 40 second delays. Significant differences are indicated by bold and italicized text.

| Group Contrast | Delay | Model<br>Estimate | 95% Lower<br>CI | 95% Upper<br>CI | p value |
| --- | --- | --- | --- | --- | --- |
| (8-15 yrs) vs. (21-29 yrs) | 2.5 | 0.62 | 0.37 | 1.03 | 0.069 |
| (16-20 yrs) vs. (21-29 yrs) | 2.5 | 1.08 | 0.55 | 2.12 | 0.959 |
| (8-15 yrs) – (16-20 yrs) | 2.5 | 1.75 | 0.86 | 3.53 | 0.152 |
| (8-15 yrs) vs. (21-29 yrs) | 5 | 0.62 | 0.38 | 1.03 | 0.068 |
| (16-20 yrs) vs. (21-29 yrs) | 5 | 1.17 | 0.60 | 2.28 | 0.844 |
| (8-15 yrs) – (16-20 yrs) | 5 | 1.88 | 0.94 | 3.75 | 0.085 |
| (8-15 yrs) vs. (21-29 yrs) | 10 | 0.87 | 0.54 | 1.42 | 0.793 |
| (16-20 yrs) vs. (21-29 yrs) | 10 | 0.76 | 0.39 | 1.49 | 0.613 |
| (8-15 yrs) – (16-20 yrs) | 10 | 0.87 | 0.44 | 1.75 | 0.893 |
| ***(8-15 yrs) vs. (21-29 yrs)*** | ***15*** | ***0.43*** | ***0.27*** | ***0.71*** | ***<0.001*** |
| (16-20 yrs) vs. (21-29 yrs) | 15 | 0.58 | 0.30 | 1.13 | 0.132 |
| (8-15 yrs) – (16-20 yrs) | 15 | 1.34 | 0.67 | 2.66 | 0.588 |
| ***(8-15 yrs) vs. (21-29 yrs)*** | ***20*** | ***0.39*** | ***0.24*** | ***0.64*** | ***<0.001*** |
| *(16-20 yrs) vs. (21-29 yrs)* | **20** | **0.39** | **0.20** | **0.76** | **0.003** |
| (8-15 yrs) – (16-20 yrs) | 20 | 0.99 | 0.49 | 1.98 | 0.999 |
| *(8-15 yrs) vs. (21-29 yrs)* | **30** | **0.42** | **0.26** | **0.69** | **<0.001** |
| *(16-20 yrs) vs. (21-29 yrs)* | **30** | **0.48** | **0.24** | **0.93** | **0.025** |
| (8-15 yrs) – (16-20 yrs) | 30 | 1.14 | 0.57 | 2.25 | 0.901 |
| *(8-15 yrs) vs. (21-29 yrs)* | **40** | **0.49** | **0.30** | **0.82** | **0.003** |
| *(16-20 yrs) vs. (21-29 yrs)* | **40** | **0.44** | **0.22** | **0.86** | **0.011** |
| (8-15 yrs) – (16-20 yrs) | 40 | 0.89 | 0.45 | 1.76 | 0.910 |
| (8-15 yrs) vs. (21-29 yrs) | 60 | 0.63 | 0.37 | 1.05 | 0.084 |
| (16-20 yrs) vs. (21-29 yrs) | 60 | 0.54 | 0.27 | 1.07 | 0.088 |
| (8-15 yrs) – (16-20 yrs) | 60 | 0.86 | 0.44 | 1.71 | 0.871 |

### Relationship Between the WFMT & DR Task

Performance in the WFMT was significantly correlated with performance in the DR task (**Fig. 3**). Significant Pearson correlations (p’s<0.028) were detected between highest and total maze in the WFMT and accuracy on the DR task for all but 2.5-s and 60-s delays. Although the correlations were nearly identical between DR accuracy and highest and total maze levels, correlations were slightly stronger between the total number of maze levels and DR accuracy for delays longer than 20 seconds.

**Fig. 3.**
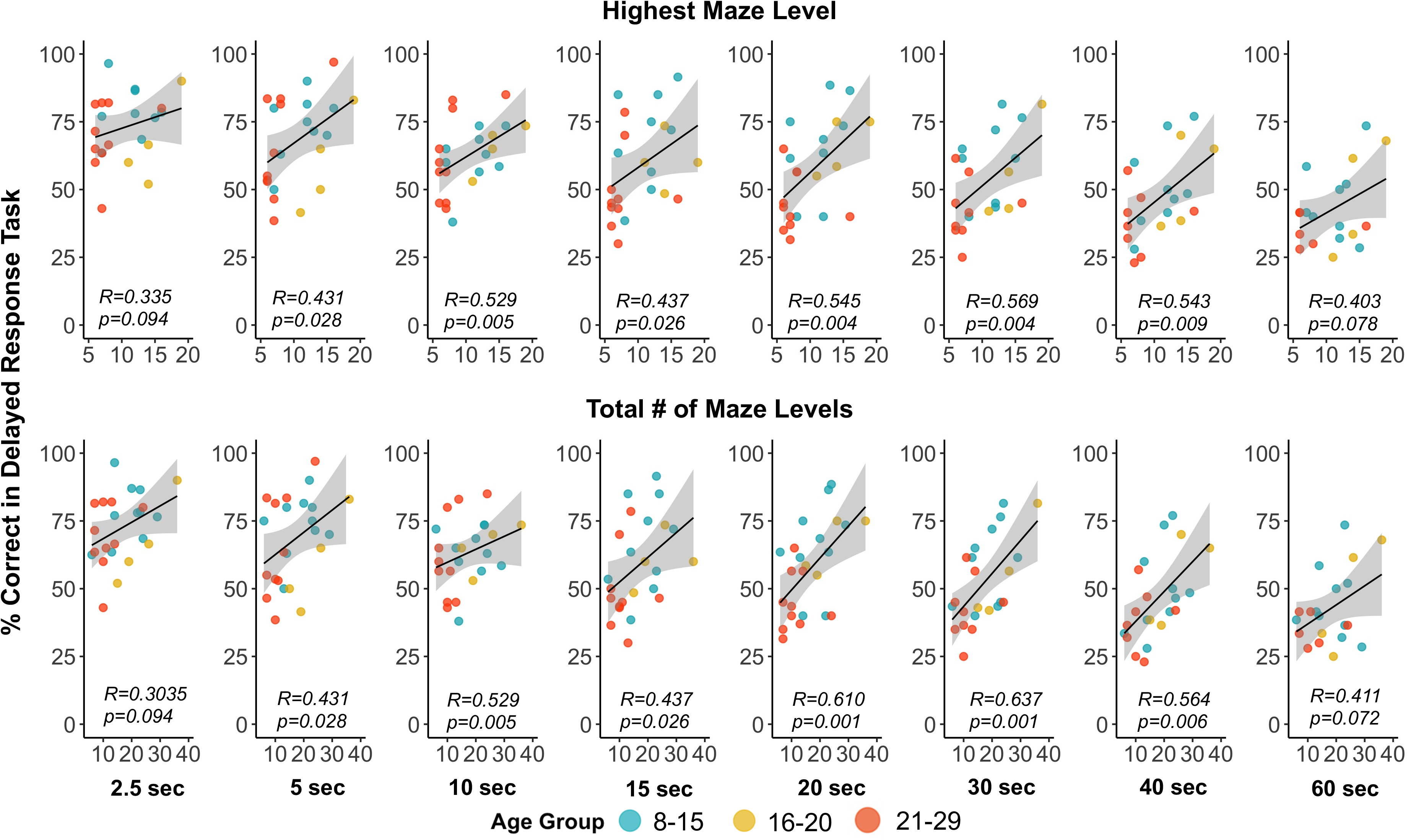
Pearson correlations between two metrics of performance in the Wake Forest Maze Task (WFMT) and accuracy (i.e., percentage correct) in the Delayed Response (DR) task of working memory by delay time. Top row: correlations between the highest maze level completed (on either the left or right series) in the WFMT and accuracy at each delay in the DR task. Bottom row: correlations between the total number of maze levels completed in the WFMT and accuracy at each delay in the DR task. Overall, performance in the WFMT agreed well with performance in the DR task (N=27)

### Gait Speed & Cognitive Performance

Gait speed declined with age; older animals exhibited slower gait speeds than younger animals (regression: t=-2.45, p=0.021). Gait speed significantly predicted performance in both the WFMT and DR task. That is, monkeys with faster gait speeds at baseline completed higher (regression: t=2.76; p=0.010; **Fig. 4A**) and more overall (regression: t=3.94; p=0.001; **Fig. 4B**) maze levels than individuals with slower gait speeds. Gait speed was also a significant predictor of the average number of successful trials at the 20 second delay (i.e., the longest delay completed by all subjects) in the DR task (regression: t=3.80; p=0.001; illustrated as percent correct in **Fig. 4C**).

**Fig. 4.**
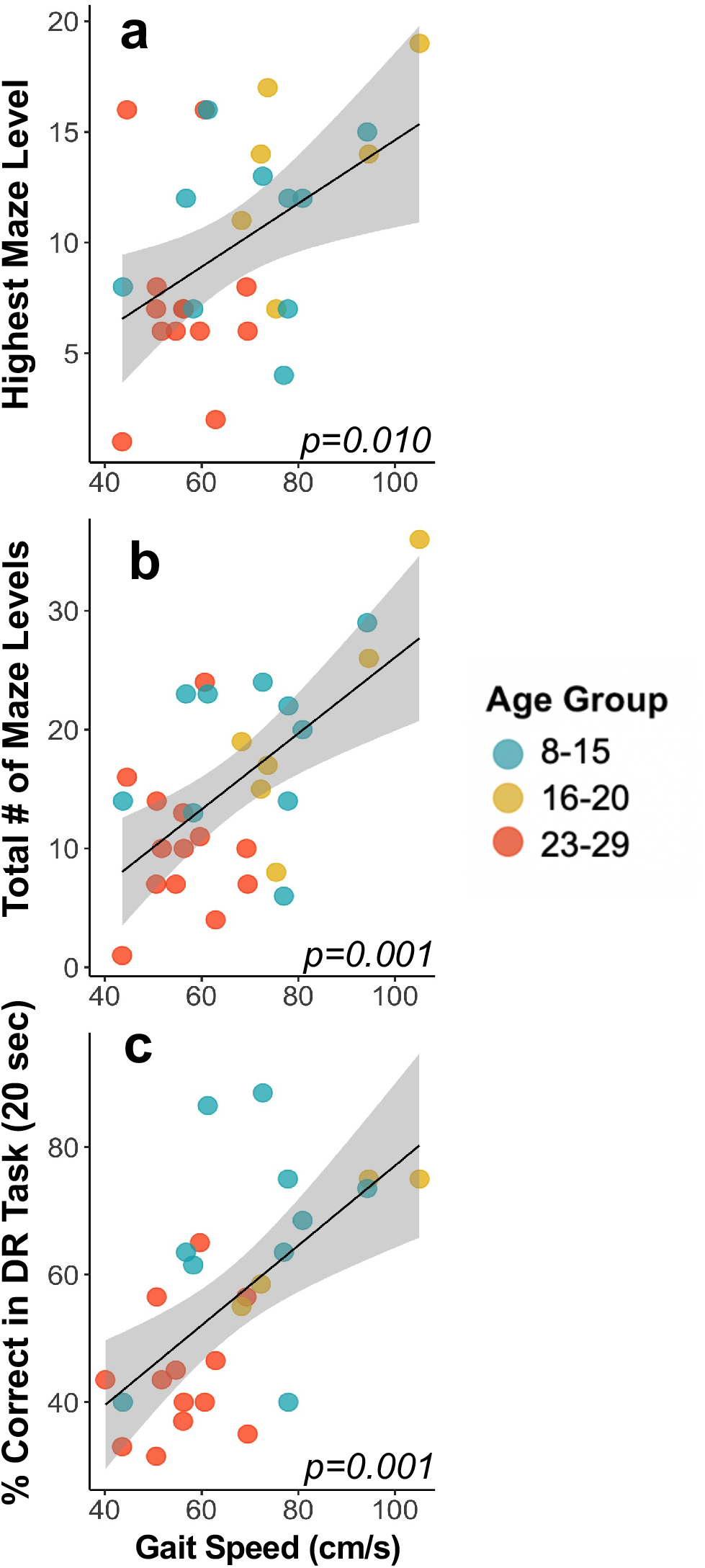
Gait speed predicted performance in the Wake Forest Maze Task and DR task. Individuals with faster gaits performed better than individuals with slower gaits. a. Highest level completed in either the left or right series (linear regression: t=2.76; p=0.010). b. Total overall levels completed (sum of left and right series) (linear regression: t=3.94; p=0.001). c. Performance, illustrated by the percent of trials correct at the 20-second delay (i.e., the longest delay reached by all subjects), in the DR task of working memory (linear regression: t=3.80; p=0.001) (a & b: N=29; c: N=27)

## DISCUSSION

To the best of our knowledge, our study is the first to simultaneously document cognitive and physical decline in nonhuman primates. Our study had three main findings. We first determined that the oldest vervets showed functional deficits in cognitive and physical performance, which adds to a growing literature espousing cognitive [36] and physical impairments [29, 37] in aged NHPs and underscores NHPs’ roles as translational models of functional decline with age. We next established that performance in the novel WFMT of executive function positively correlated with performance in the well-established DR task of working memory. Finally, we demonstrated an association between gait speed and cognitive performance at one-year follow-up, such that slower gait speeds predicted poorer performance in both cognitive domains. Taken together, these findings suggest that vervets exhibit age-related functional impairments in ways similar to humans [15, 16, 20, 17–19] and that slow gait may presage cognitive decline.

Performance in the WFMT agreed well with performance in the DR task of working memory. The WFMT has several advantages over other experimental tools. The WFMT protocol requires brief periods of social separation, approximately 30 minutes per day, in which individuals could remain within their home environment. Separation from social groups reliably induces stress responses in NHPs [38–40], which impacts learning and cognitive performance [41–43]. Reducing durations of social separation also minimizes disruption to group dynamics. This is important for study validity because social factors directly influence NHP behavior and health [44, 45, 38], which may confound results. The WFMT also considerably reduces time required for training and testing. The majority of our sample (27 of 29) completed the WFMT protocol over the course of two weeks (i.e., 5 experimental hours), whereas the DR task required an average of 21 days (i.e., 9-17 experimental hours). Furthermore, age did not impact the rate with which animals acquired the cognitive tasks, which suggests that inter-individual differences in motivation or perception were unrelated to age. Notably, the study subjects had not recently undergone any cognitive testing paradigms before the WFMT but not the DR task. The ability to gather valid data in a timely manner, with minimal stress, eliminates many key barriers to assessing cognitive capabilities in NHPs.

In addition, the WFMT battery may reflect function across a broad range of cognitive capabilities – e.g., problem solving, behavioral flexibility, self-control, and planning – instead of focusing on performance within a single domain. The capacity to measure complex processes associated with executive function may bolster the WFMT’s translational value. Clinical assessments executive function (e.g., Wisconsin Card Sorting Test, Stroop Task, Go/No-Go Task, and Flanker Task) are important paradigms for diagnosis of mild cognitive impairment [46] and AD [47], and deficits in executive function may be some of the earliest cognitive domains impacted by AD [48]. Thus, the WFMT may be particularly well-suited for preclinical, longitudinal studies of neurodegenerative disorders. Notwithstanding these potential strengths, this proposal requires further studies of construct and predictive validity. For example, subsequent research is needed to determine whether performance in the WFMT accurately discriminates neuropathological profiles.

Gait speed was significantly associated with cognitive performance at one year in both the WFMT and the DR task. This period of time translates to roughly a three-to-four year follow up in humans, based on the maximum lifespans of vervets (~30 years [49]) and humans (~122 years [50]). The association between slow gait and later cognitive impairment aligns well with longitudinal clinical surveys showing similar temporal dynamics between physical and cognitive impairment (e.g., [15, 16, 20, 17–19]). Slow gait may represent an early marker of progressive cognitive dysfunction, and progressively slowed gait and cognitive impairment may result from a shared neuropathology, as has been suggested in humans [6]. Early detection of neuropathology and cognitive impairment may provide important information about individuals’ risk, well before cognitive symptoms emerge. Ultimately, this information may provide a straightforward means of screening in large NHP populations that is not possible with formal cognitive testing, and with advances in automatic motion capture it could be possible to acquire data on individual trajectories of gait speed in large populations.

While our study had several important strengths, its limitations should be considered. Aging females constitute our study cohort. Husbandry at the VRC recapitulates the normal social organization of wild vervets, in which the adult sex ratio is approximately 1:5. The availability of older-aged male monkeys therefore is limited. Another potential limitation is that our cohort consists of both reproductively active and reproductively senescent females. Additional study of the effects of reproductive status on cognitive performance within age groups would provide important insights, but such an investigation was beyond the scope of this project. Importantly, we were limited to a single physical and cognitive assessment per individual. Repeated measures would provide information needed to fully characterize cognitive performance over time and enable analyses of the implications of different rates of change for individuals’ health. In addition, longitudinal data need to be collected in other cohorts to fully characterize the natural history of cognitive and physical decline in NHPs. Future investigations will examine the relationships between physical and cognitive performance and other AD biomarkers – e.g., neuroanatomy, cerebrospinal fluid biomarkers, and histopathology. Such multidimensional approaches will bolster a more holistic understanding of brain-body connections and illustrate the functional implications of neuropathological phenotypes across the lifespan.

## Conclusions

In conclusion, we have provided important information that advances vervet monkeys as translationally relevant models of physical and cognitive decline. Moreover, our data suggests that slow gait may represent a risk factor for cognitive decline. However, this hypothesis needs to be confirmed with additional physical and cognitive assessments. Gait speed may represent a simple, inexpensive biomarker of cognitive performance. This information is critical for informing future interventions which aim to detect and target modifiable risk factors before symptoms manifest. This study helps to clarify the patterns typifying systemic age-related functional declines and may ultimately be used to inspire new interventions to prevent and treat the progression of age-related cognitive impairment and Alzheimer’s disease.

## ACKNOWLEDGMENTS

We thank the Chrissy Long and Justin Herr for their advice and assistance during this study. The content is solely the responsibility of the authors and does not necessarily represent the official views of the NIH.

## SOURCES OF FUNDING

This work was supported by several mechanisms, including: National Institutes of Health (NIH) R01HL087103 (CAS), NIH RF1AG058829 (CAS & SC), P30 AG049638 (SC), Intramural Grant from the Department of Pathology, Wake Forest School of Medicine (CAS), Wake Forest Claude Pepper Older Americans Independence Center grant P30 AG21332 (SK), Vervet Research Colony (P40-OD010965) (MJ), and the Wake Forest Clinical and Translational Science Institute (NCATS UL1TR001420).

## DISCLOSURES

The authors have no conflicts of interest to disclose.

## Notes

### Competing Interest Statement

The authors have declared no competing interest.

