## Supplemental Information for "Temporal emergence of age-associated changes in cognitive and physical function in vervets (*Chlorocebus aethiops sabaeus*)"

Journal: GeroScience

^1^ Department of Pathology/Comparative Medicine, Wake Forest School of Medicine

^2^ Sticht Center for Healthy Aging and Alzheimer’s Prevention, Department of Internal Medicine - Gerontology and Geriatric Medicine, Wake Forest School of Medicine

^3^ Wake Forest Alzheimer’s Disease Research Center

^4^ Nash Family Department of Neuroscience, Icahn School of Medicine at Mount Sinai

Corresponding Author:	Carol A. Shively, PhD

Department of Comparative Medicine/Pathology

Wake Forest School of Medicine

Medical Center Blvd

Winston-Salem, NC 27157-1040

**SUPPLEMENTARY INFORMATION**

**Supplementary Tables**

**Supplementary Table 1** Study subject characteristics.

| **Age Group (years)** | **Avg. Age** | **N_Gait_** | **N_WFMT_** | **N_DR_** | **Avg.**  **Gait ± SE**  **(cm/s)** |  | **Avg. Highest Maze Level** | **Avg. Total**  **Maze Levels** | **Reached Longest Delay (%)** |
| --- | --- | --- | --- | --- | --- | --- | --- | --- | --- |
| 8-15 | 11.16 | 10 | 10 | 10 | 71.22±4.39 |  | 10.6 (range: 4-16) | 18.8 (range: 6-29) | 100% |
| 16-20 | 19.69 | 6 | 6 | 4 | 81.55±6.06 |  | 13.7 (range: 7-19) | 20.2 (range: 8-36) | 100% |
| 21-29 | 24.51 | 14 | 13 | 13 | 55.02±2.40 |  | 7.4 (range: 1-19) | 10.3 (range: 1-24) | 46% |

**Supplementary Figures**

**Supplementary Fig. 1** Wake Forest Maze Task (WFMT) configuration sequence from acclimation to testing. Acclimation involved 1) training the monkey to interact with the puzzle feeder by loading a randomly assorted maze with high quality food rewards (grapes) (Phase 01); 2) replacing high quality foods with less desirable (but more durable) food items (carrots) followed by positive reinforcement (i.e., clicker mark followed by the high-quality reward) (Phase 02); and 3) practice in moving a food reward over a vertical insert (Phase 03). Following Acclimation, monkeys were tested using increasingly difficult maze configurations. Pilot data indicated a directional preference across individuals, so we created left and right mirror images for each level. Thick blue lines indicate removable tabs; the orange rectangle represents the food reward (carrot); and the ovals represent holes through which the monkey could manipulate the reward. The reward could only be retrieved through the enlarged openings (illustrated by the rounded rectangles) at the bottom of the maze.

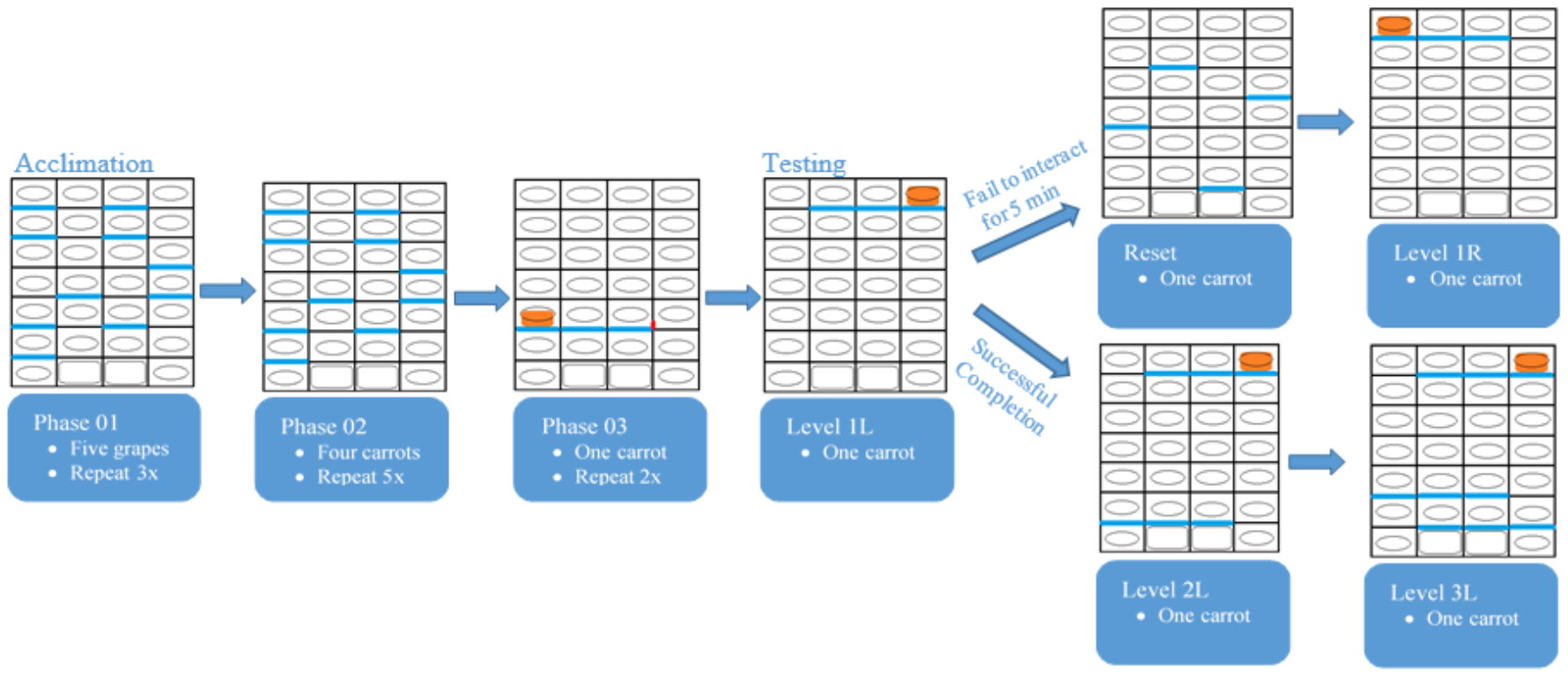

**Supplementary Fig. 2** Wake Forest Maze Task (WFMT) levels 1-19. Each level consisted of left and a right mirror image. “L” designates the left series and “R” designates the right series.

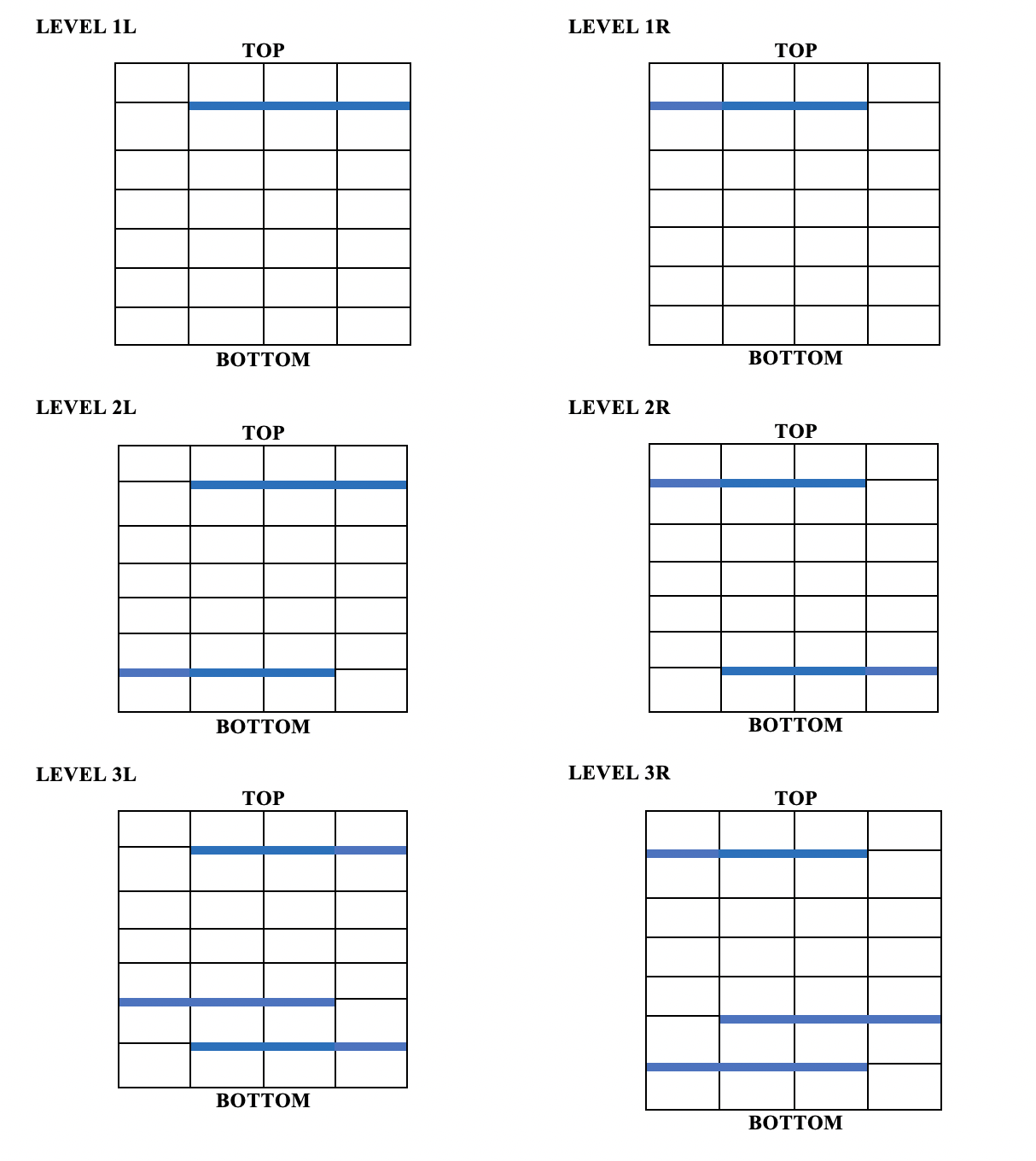

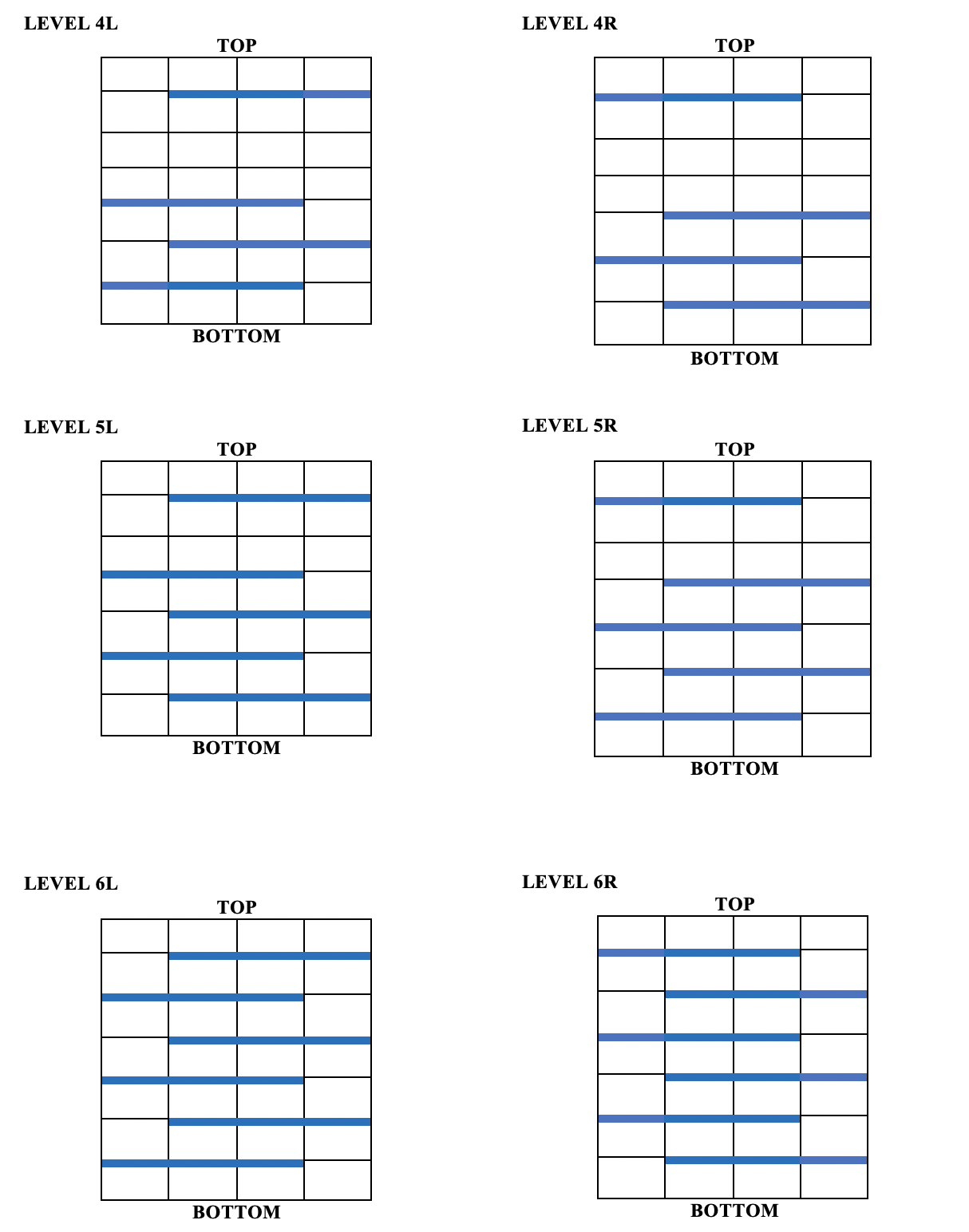

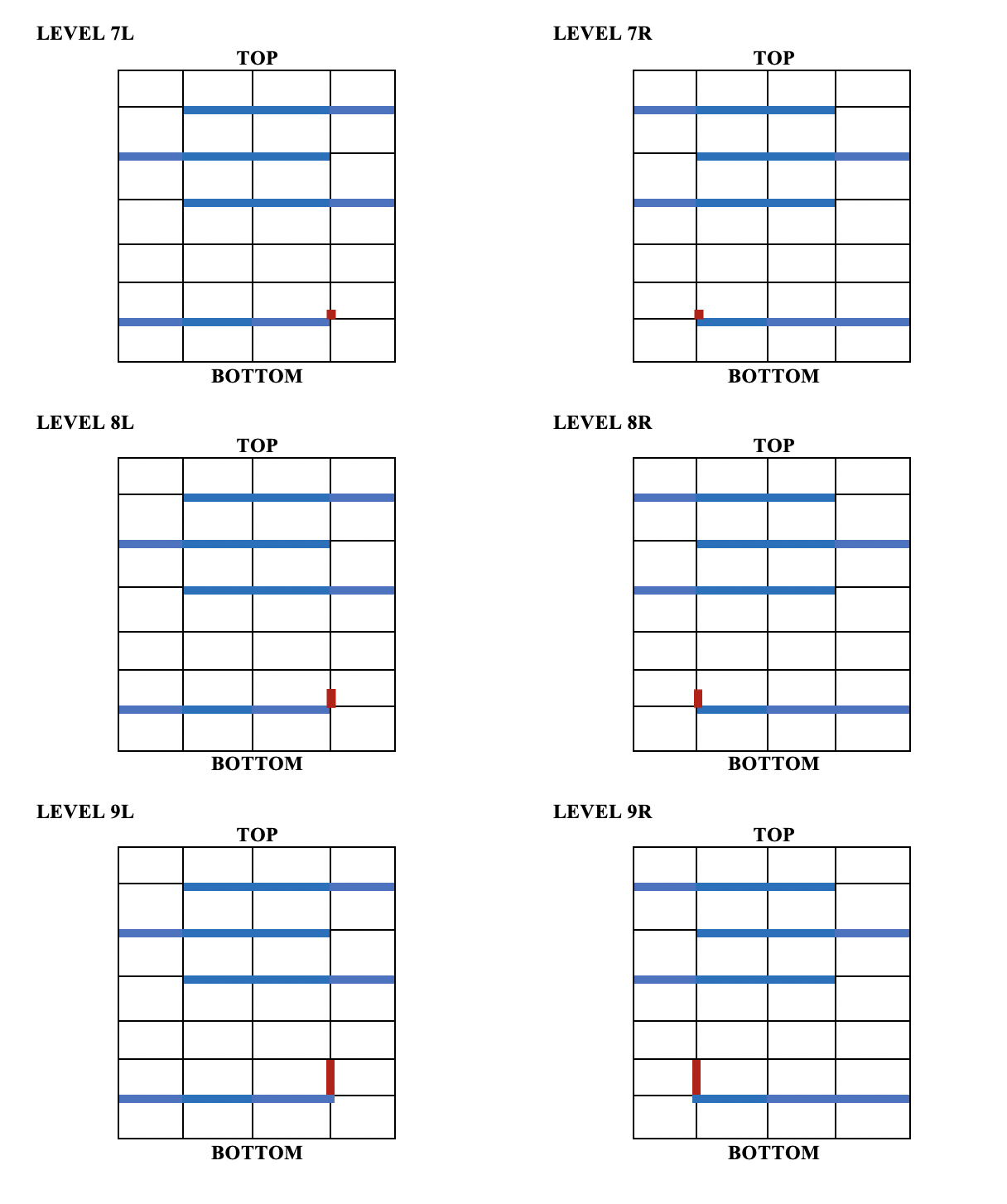

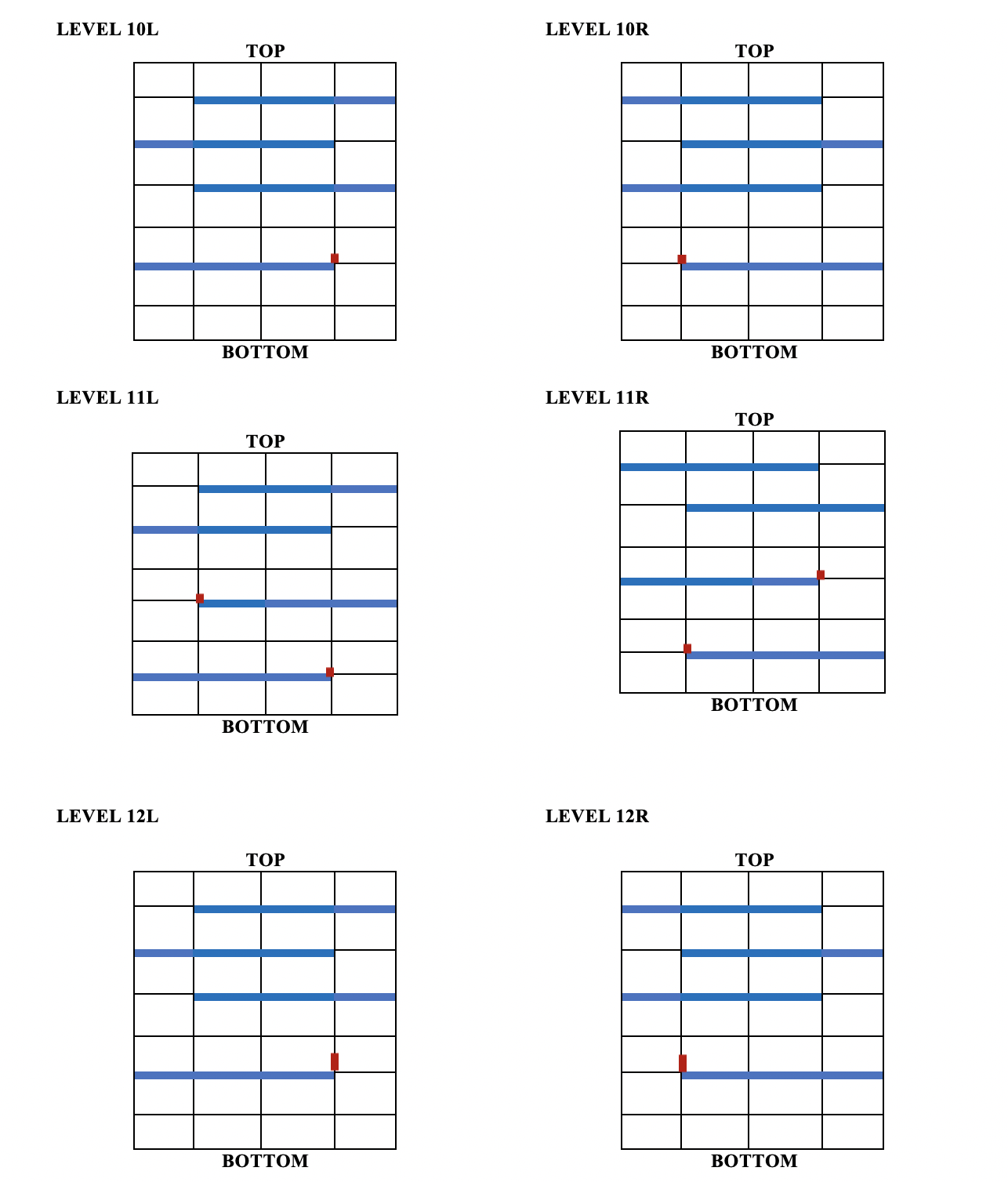

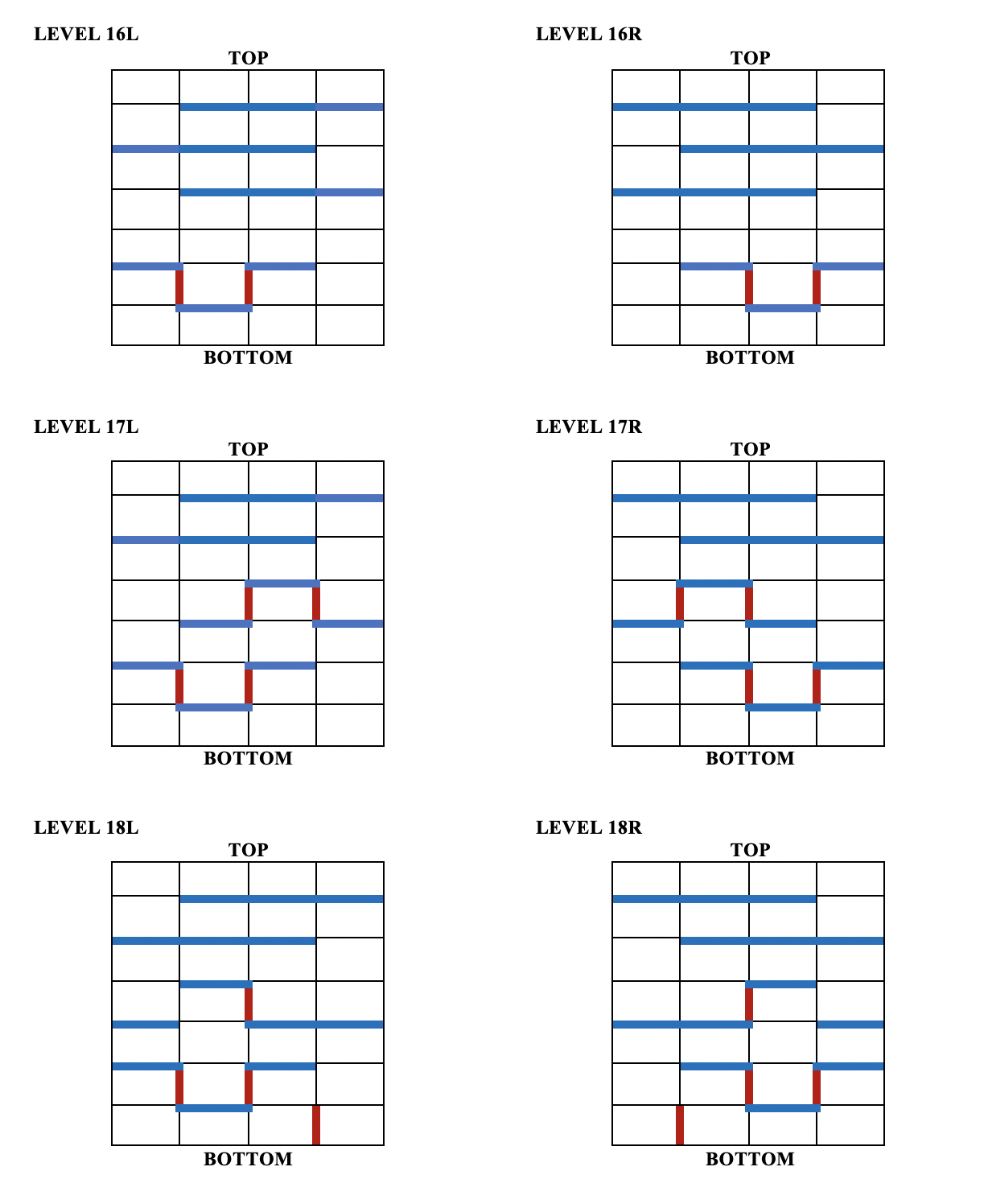

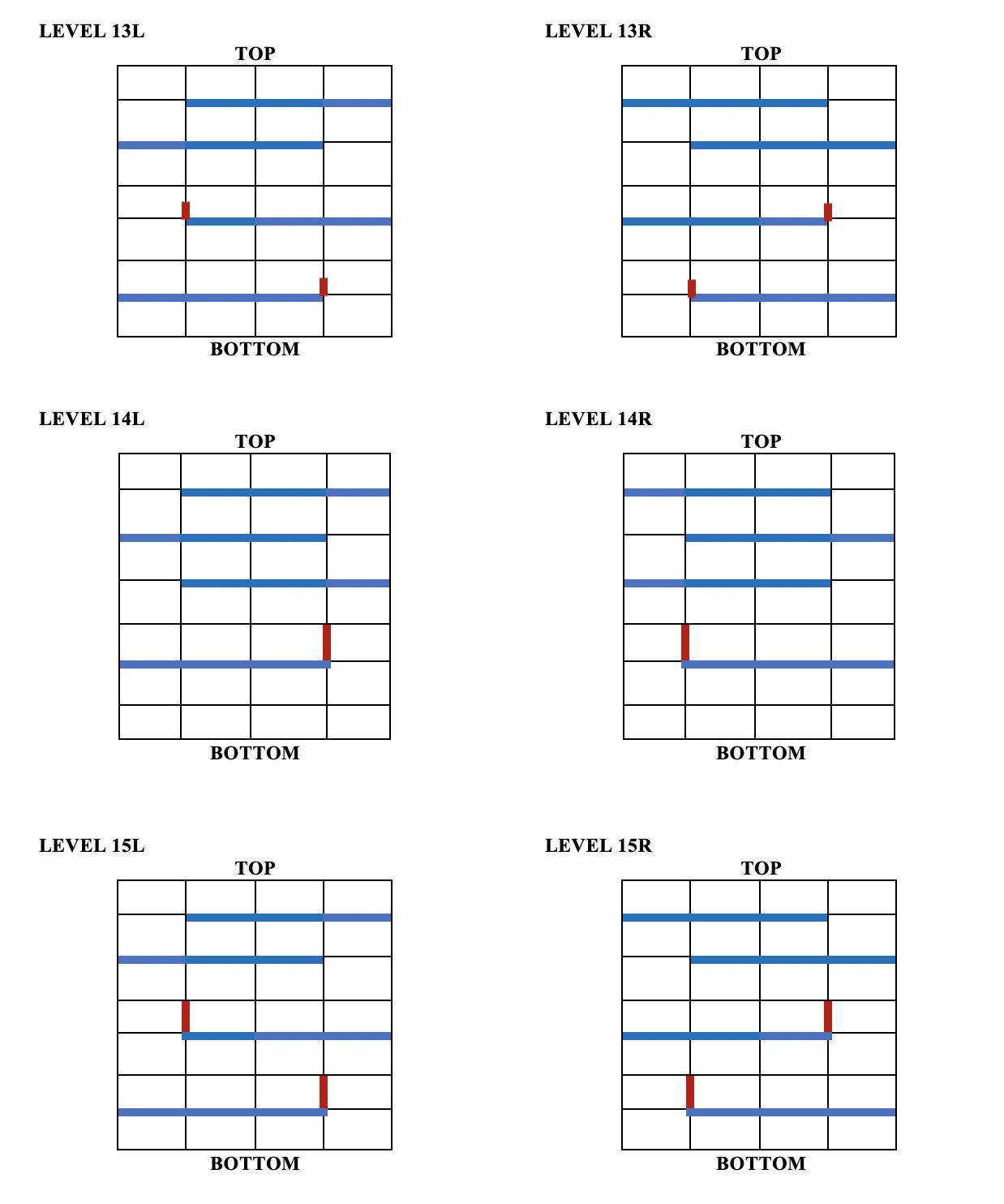

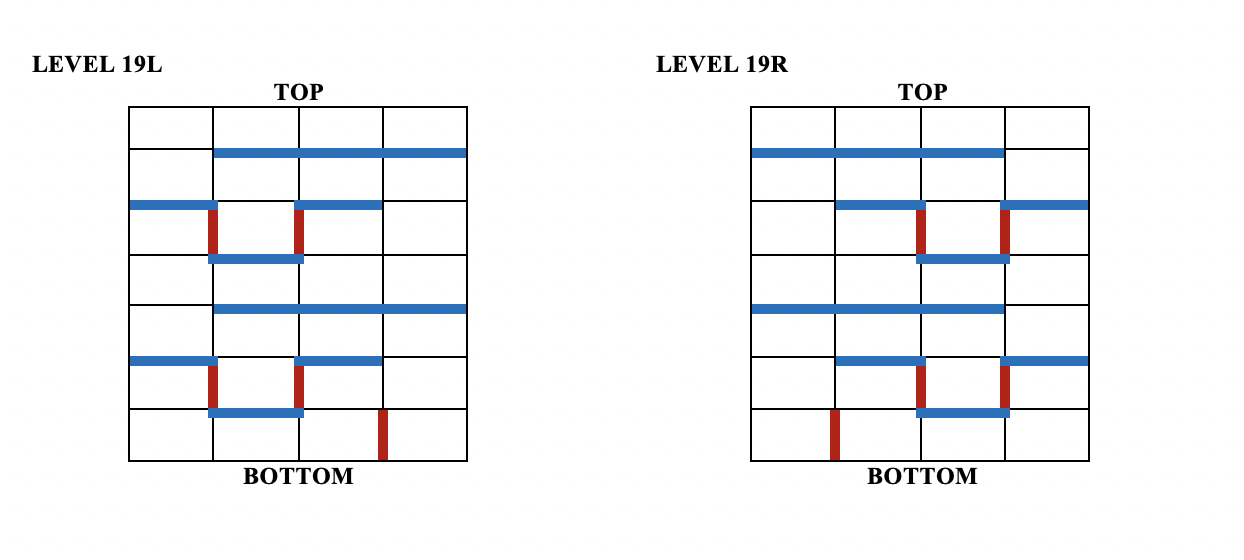

**Supplementary Fig. 3** The Delayed Response (DR) task was modified from a Wisconsin General Test apparatus to assess working memory. A) The apparatus consisted of three opaque boxes which the subject could access when a plexiglass screen was raised. B) Food rewards were randomly placed in one of three opaque boxes. The red dot indicates the baited box. If the monkey interacted with the baited box first, the response was recorded as correct (“+”). If either empty box was chosen first, the response was recorded as incorrect. (“-”). If there was no interaction with the apparatus for 30s, the trial was scored as omission.

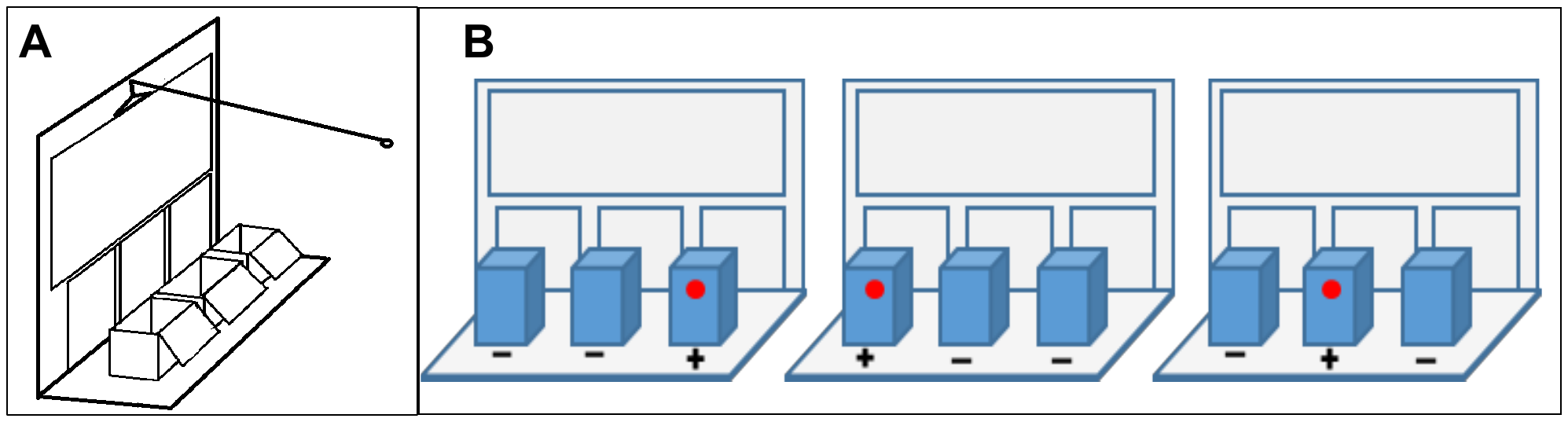

**Supplementary Fig. 4** Number of animals to reach the longest delay (60 seconds) in the Delayed Response (DR) task of working memory differed across age groups (χ^2^=10.177, df=2, p=0.006). All middle and older-aged vervets reached the longest delays, whereas the majority of oldest vervets failed to reach the longest, i.e., most difficult, delay.

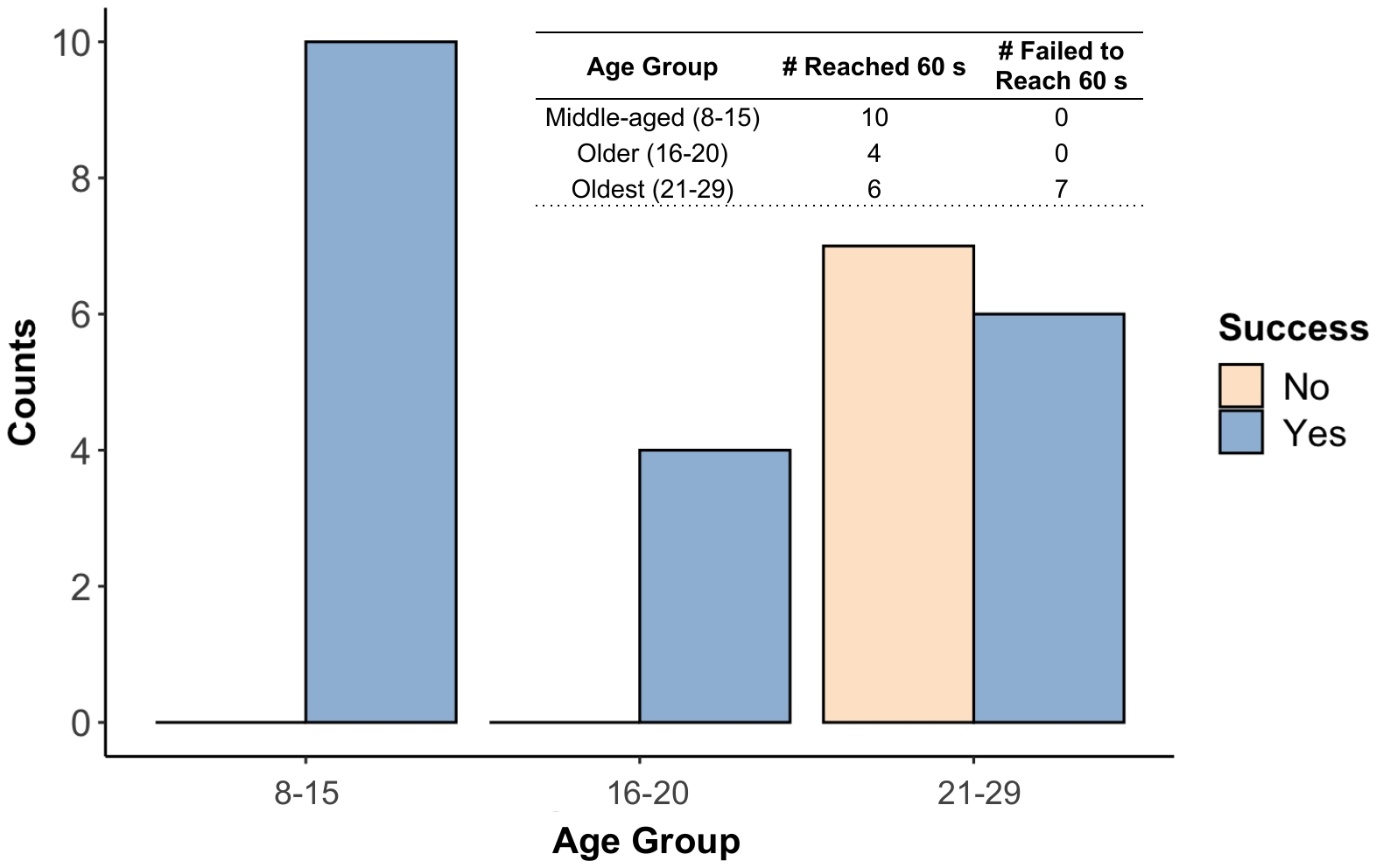
